# Visualization of SARS-CoV-2 Infection Scenes by ‘Zero-Shot’ Enhancements of Electron Microscopy Images

**DOI:** 10.1101/2021.02.25.432265

**Authors:** Jakob Drefs, Sebastian Salwig, Jörg Lücke

**Affiliations:** Machine Learning Lab, School of Medicine and Health Science Carl von Ossietzky University Oldenburg, Germany

## Abstract

Electron microscopy (EM) recordings of infected tissues serve to diagnose a disease, and they can contribute to our understanding of infection processes. Consequently, a large number of EM images of the interaction of SARS-CoV-2 viruses with cells have been made available by numerous labs. However, due to EM recording techniques at high resolution, images of infection scenes are very noisy and they appear two dimensional (‘flat’). Current research consequently aims (A) at methods that can remove noise, and (B) at techniques that allow for recovering a 3D impression of the virus or its parts. Here we discuss a novel method which can recover a spatial impression of a whole infection scene at high resolution. In contrast to previous approaches which aim at the reconstruction of single spike proteins or a single virus, the here used method can be applied to a single noisy EM image of an infection scene. As one example image, we show a high resolution image of SARS-CoV-2 viruses in Vero cell cultures (Fig. 1). The method we use is based on probabilistic machine learning algorithms which can operate in a ‘zero-shot’ setting, i.e., in a setting when just one noisy image (and no large and clean image corpus) is available. The probabilistic method we apply can estimate non-noisy images by inferring first order statistics (pixel means) across image patches using a previously learned probabilistic image representation. Estimating higher order statistics and appropriately chosen probabilistic models then allow for the generation of images that enhance details and give a spatial impression of a full nanoscopic scene.

## Introduction

The SARS-CoV-2 virus which causes the COVID-19 disease is currently infecting a high percentage of humans world-wide, which has dramatically impacted society and economy. Electron microscopy (EM) imaging is an established method to visualize viruses and infection processes. Such images can contribute to our understanding of how a given virus infects a cell, how it binds to cells and different tissues, and how it spreads in the body (Lamers et al., 2020; Ivanova et al., 2016, for two examples). Transmission electron microscopy (TEM) allows for recordings of high-resolution images on the scale of a few nanometers. However, for such high-resolution / low-light conditions, EM images contain significant amounts of noise. Furthermore, images of viruses and infection processes appear two dimensional (‘flat’) at this resolution (Fig. 2, top-left). In order to address the problem of high noise, algorithms for noise removal have become a very established tool, and they can significantly improve further image processing as well as the interpretation of images by humans. For images of SARS-CoV-2 viruses, a further goal has been the reconstruction of the virus or virus parts in 3D from EM data. On the most detailed level, work e.g. by Wrapp et al. (2020); Ke et al. (2020) has used large amounts of TEM images of the SARS-CoV-2 particles in order to reconstruct the protein’s 3D shape on a molecular level. It was thus possible to visualize a 3D model of the protein in different conditions which can help to better understand the protein binding process. On the next lower resolution level, very recent work by Nanographics GmbH has released a 3D image of the SARS-CoV-2 virus using cryo-electron tomography (Nanographics, 2021; Yao et al., 2020). The approach is based on a large number of TEM images of the same virus recorded as the sample is tilted along an axis. These images are then merged to a 3D image using technology developed for computer tomography (compare, e.g., Doerr, 2017).

**Figure 1:**
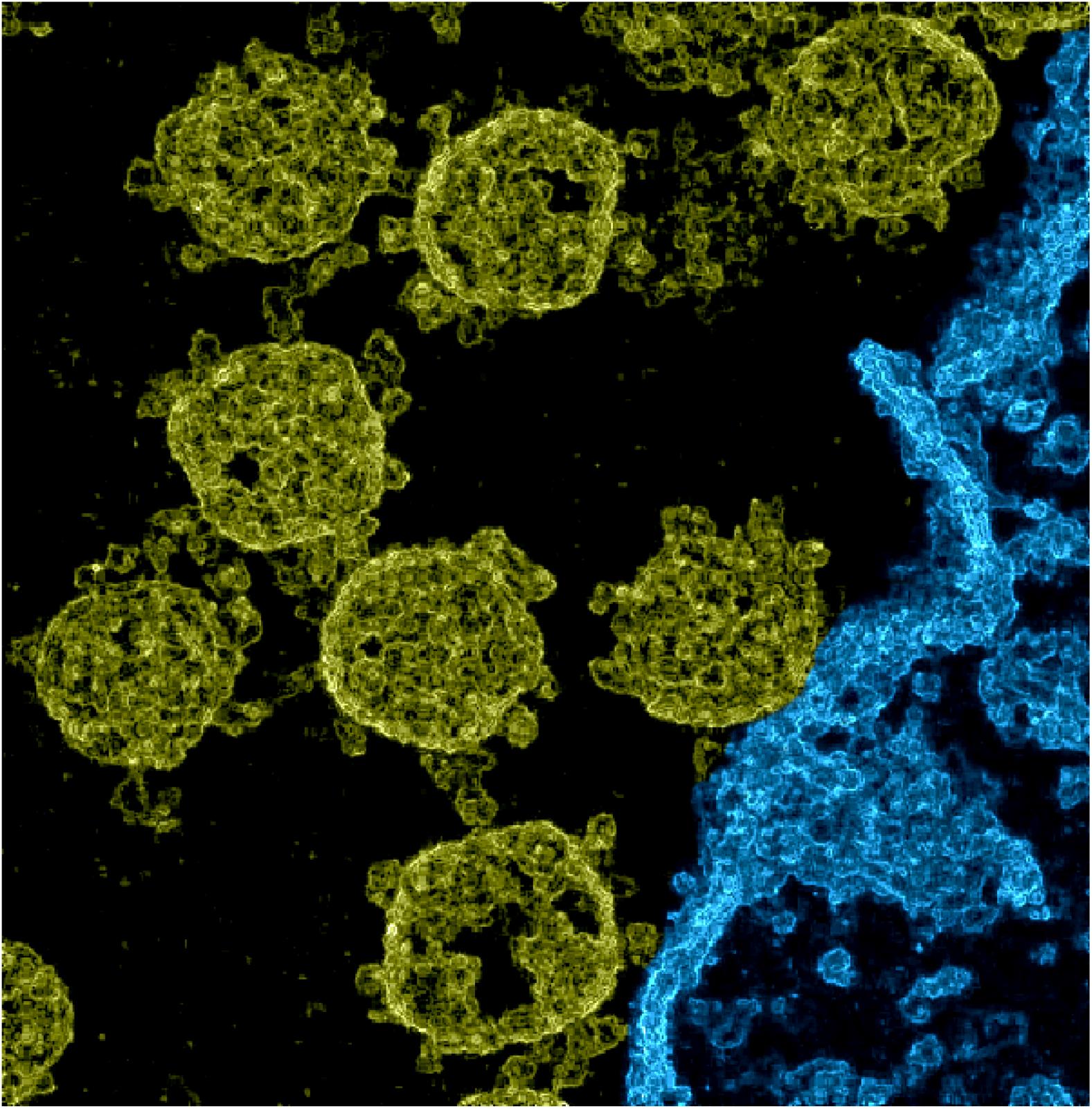
Image of SARS-CoV-2 viruses in Vero cell cultures. We used data made available by Laue et al. (2021) who used transmission electron microscopy of ultrathin plastic sections. Based on the data, we estimated pixel variances during the application of probabilistic Machine Learning algorithms for denoising (Sheikh et al., 2014; Monk et al., 2018; Drefs et al., 2020). The resulting image of these variances gives a spatial impression (also see Supplementary Figure 1, which shows a close comparison between a section of the noisy EM image and the corresponding estimated pixel variances). The here shown image was then obtained after contrast enhancement and colorization: structures that we manually identified as belonging to a cell were colored in blue (lower right area), the remainder was colorized in yellow. The image without any manual colorization or contrast enhancement is shown as Fig. 2 (top right) of this document. The colored and the original image are available for download at this repository.

**Figure 2:**
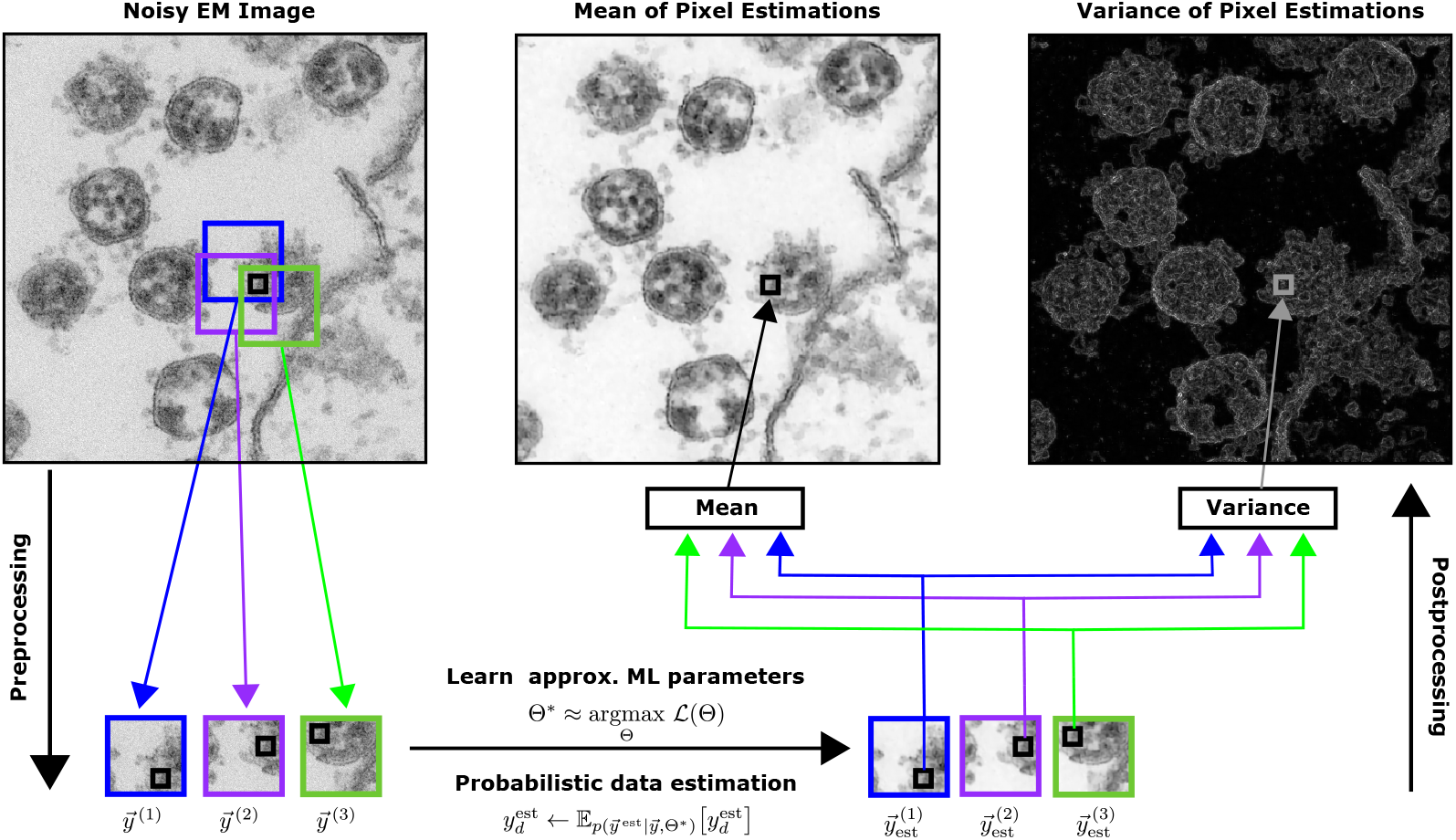
‘Zero-Shot’ Enhancement of an Electron Microscopy Image of SARS-CoV-2 viruses using probabilistic Machine Learning algorithms for denoising. The input (top left) is a cropped and scaled version of the freely available file `Dataset 07 SARS-CoV-2 077.tif` from Laue et al. (2020, 2021). Using patches cut out of the noisy input, we learn a probabilistic representation of the EM image using a linear spike-and-slab sparse coding model. We can then apply the learned representation to probabilistically reconstruct each EM image patch. Finally, we generate reconstructed EM images (top center and right) by computing means and variances of the pixel estimates from different patches. The EM image enhancement approach does not require (clean) training data and can directly be applied to a single noisy image (see text for details).

Here, we consider a method that can operate on a single TEM image, i.e., a method that neither requires the processing of a large number of different images of different spike proteins (such as Wrapp et al., 2020; Ke et al., 2020) nor the processing of many images of the same virus (as Nanographics, 2021). Instead, we consider a method that can be applied to a single noisy TEM image in order to enhance the image. We use so-called probabilistic generative models to first learn an image representation. If only the noisy image itself is available for learning, the image processing task is commonly referred to as a ‘zero-shot’ task (compare, e.g., Shocher et al., 2018; Chen et al., 2020; Soh et al., 2020). The ‘zero-shot’ setting is often considered as the most challenging setting. Representations based on single images often have the advantage, however, that the intrinsic image statistics contains higher quality information than statistics learned from large image corpora (e.g. Shocher et al., 2018, for a recent discussion). Once our method has learned a representation for a noisy TEM image of a nano-scale infection scene, we use this representation for two tasks: (A) for standard image denoising, and (B) for the generation of images which can recover a spatial impression of the originally ‘flat’ appearing image.

## Results

As main example image for our method we used a freely available transmission electron microscopy (TEM) image of SARS-CoV-2 viruses in Vero cell cultures (Laue et al., 2020, 2021). The original image was recorded with 1376 x 1032 pixels with a pixel edge length corresponding to 0.54 nm. From this image, we cropped a section of 1016 × 1032 pixels containing SARS-CoV-2 viruses and parts of a Vero cell. In a further preprocessing step we scaled this cropped image to a size of 508 × 516 by omitting every second pixel from the width and the height of the image. This interpolation step was performed in order to reduce the computational demand and at the same time not to change the image statistics. Fig. 2 (top left) shows this image.

Fig. 2 (top middle) shows the result of a ‘zero-shot’ denoising algorithm applied to the original image of Fig. 2 (top left). We used a linear generative model with spike-and-slab prior (Yoshida and West, 2010; Goodfellow et al., 2013; Sheikh et al., 2014; Drefs et al., 2020) which learns a representation from 6 × 6 patches of the noisy TEM image. The model can then infer the most likely underling non-noisy image. Training and inference is based on recent truncated variational optimization (Guiraud et al., 2018; Drefs et al., 2020) with details given in the Methods section below. To generate the denoised image, our approach estimates the mean of each pixel (i.e., statistically the first moment). If we instead use the learned representation to compute the variance of each pixel’s mean estimation (i.e., a higher order reconstruction statistics), we obtain the image shown in Fig. 2 (top right). Displaying the variance of pixel estimation removes the direct information on matter density from the TEM image. Information about boundaries between high and low density matter is maintained because at boundaries, image shapes vary more strongly than in volumes with uniform matter distribution. The visualization of pixel variance based on a sufficiently sophisticated statistical model of image patches in Fig. 2 (top right) recovers a spatial impression of the SARS-CoV-2 viruses and of the whole infection scene. The introductory figure (Fig. 1) is based on Fig. 2 (top right) but was manually colored (structures that we manually identified as belonging to the cell were colored in blue, the remainder was colorized in yellow).

The same image we also processed using alternative probabilistic models. For instance, Fig. 3 uses a Gamma-Poisson mixture model (Monk et al., 2018) instead of the spike-and-slab linear model used for Figs. 1–2 (see Methods and caption of Fig. 3 for details). As can be observed, the model itself changes the spatial appearance and (to a lesser extend) also the appearance of the conventionally denoised image (compare Fig. 3 (left) with Fig. 2 (top middle)). Also the hyper-parameters which are used for the respective model change denoised image and corresponding spatial representation (see Fig. 4 for examples and Methods for details). We found that models which provide an explicit probabilistic distribution for image patches are important for our purposes, while, for instance, non-local image processing methods such as the BM3D baseline are less suitable, particularly when hyperparameters that require a priori knowledge, e.g about the data noise level, cannot be set optimally (see Fig. 6).

**Figure 3:**
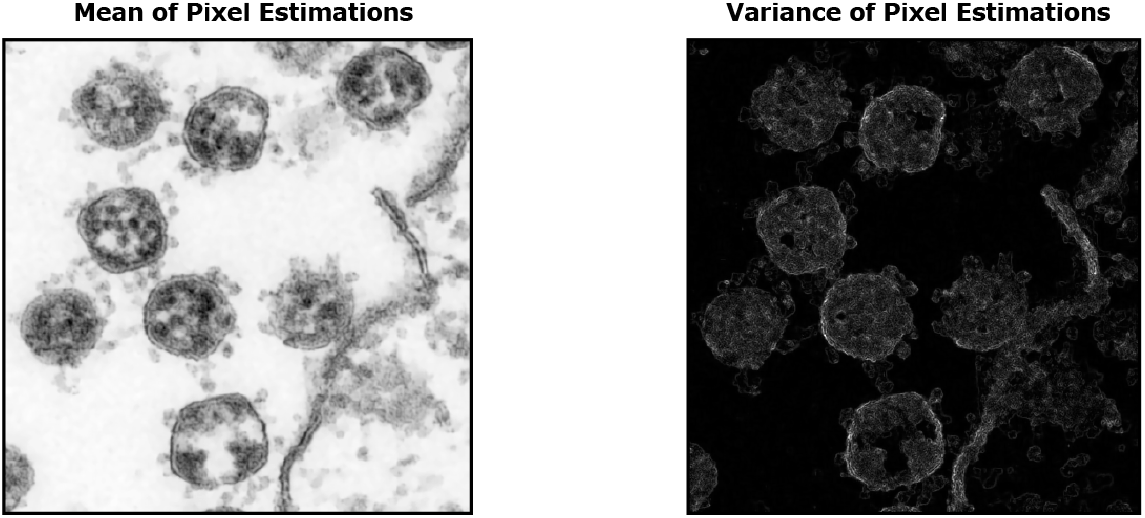
The corresponding results as shown in Fig. 2 (top center and right) when applying a Gamma-Poisson Mixture model to the noisy EM image from Fig. 2, top left (see text for details).

**Figure 4:**
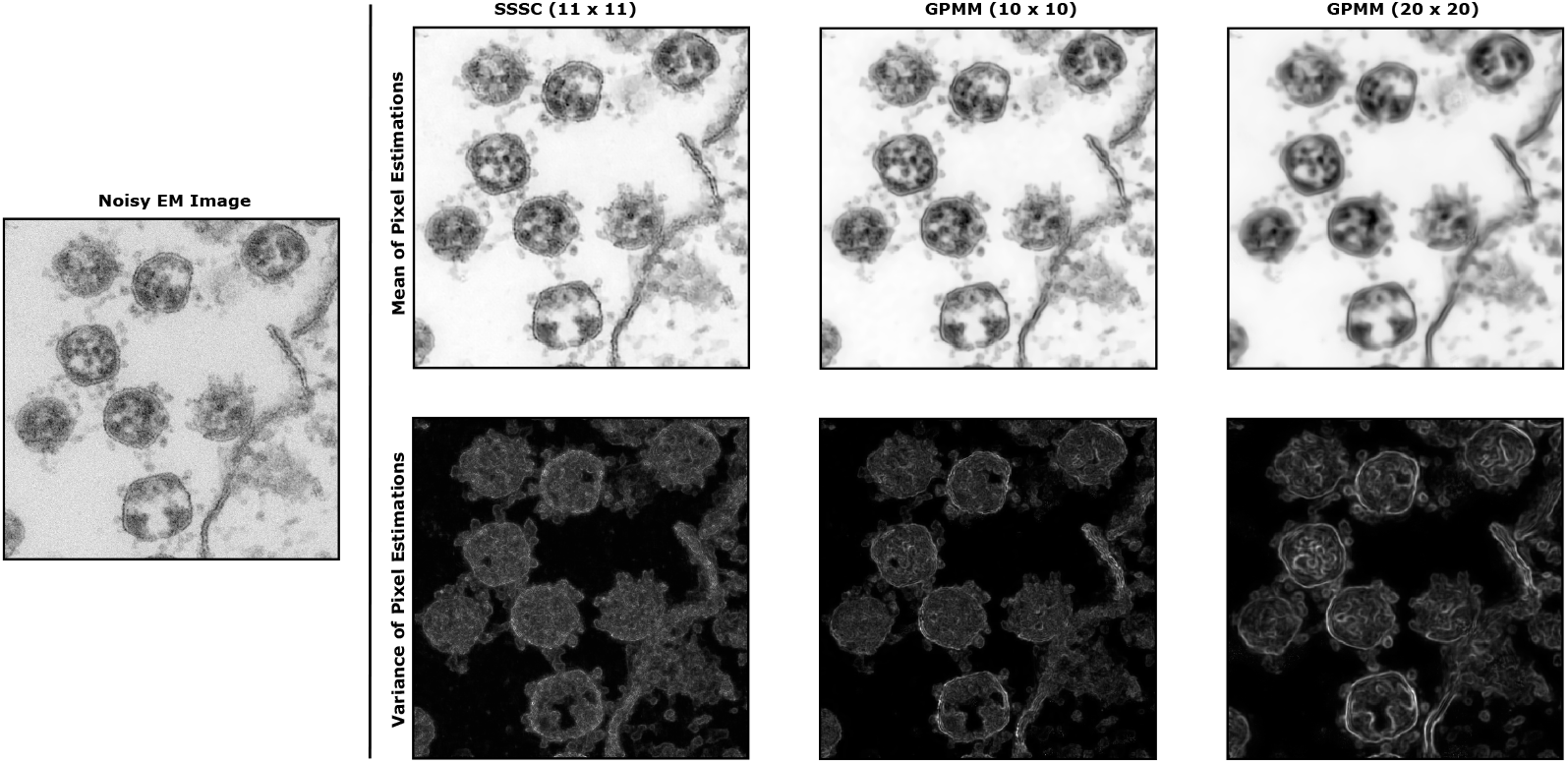
‘Zero-Shot’ Enhancements of the noisy EM image from Fig. 2 (top left) obtained with SSSC models and GPMMs using different patch sizes than in Figs. 2 (top center and right) and 3, respectively.

We provide further results with different models and settings and for different images: Fig. 5 shows, for instance, a part of Fig. 3 with variance reconstruction using a GPMM model. Fig. 7 shows an image of SARS-CoV-1 viruses (the virus that caused the outbreak of the SARS disease in 2002/2003).

**Figure 5:**
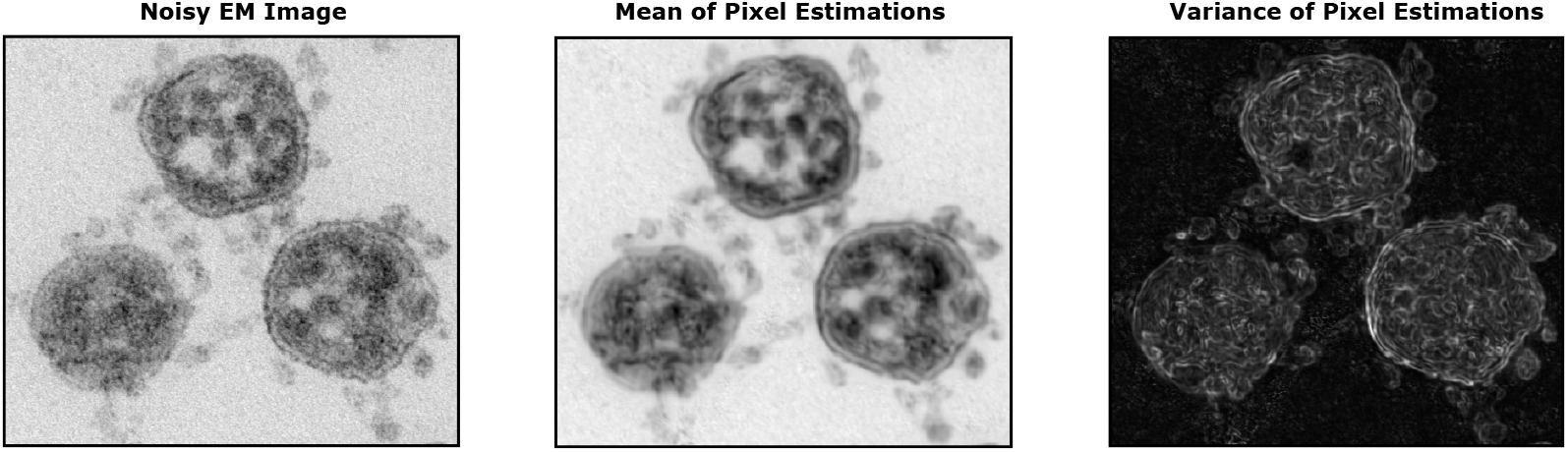
‘Zero-Shot’ Enhancements of an unscaled 523 × 471 crop of the image `Dataset 07 SARS-CoV-2 077.tif` from Laue et al. (2020). Results obtained with a GPMM trained on 20 × 20 EM image patches (see Methods for details).

**Figure 6:**
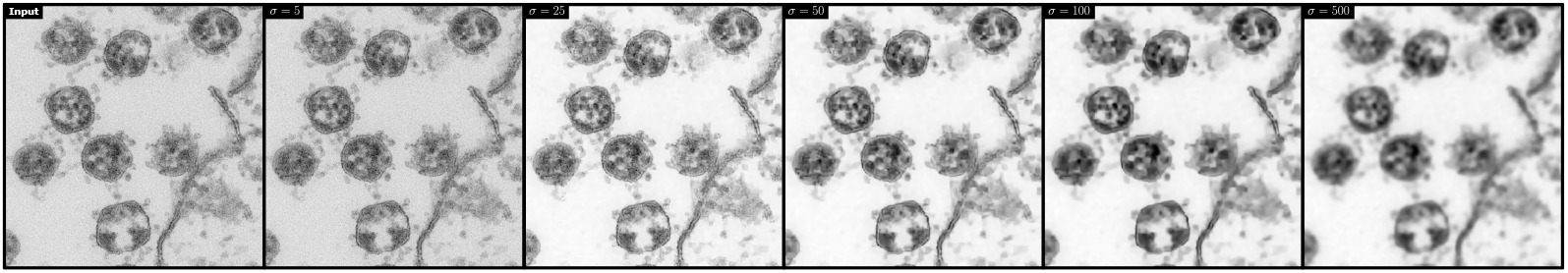
Denoised output of a noisy EM image using the BM3D baseline with hyperparameters *σ* = 5, 25, 50, 100, 500. The input image (left) is the same as in Fig. 2 (top left).

**Figure 7:**
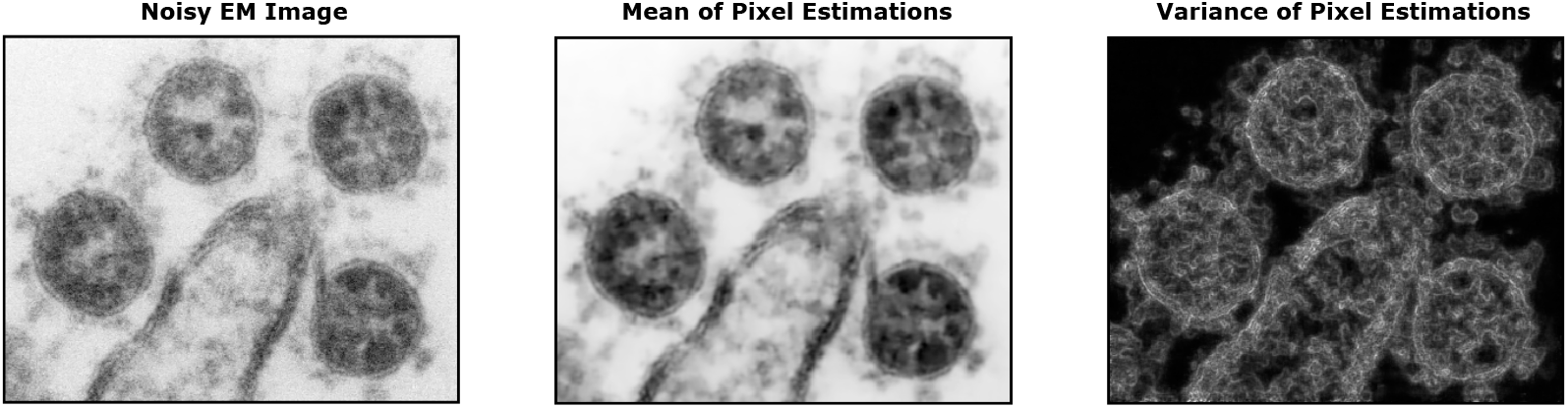
‘Zero-Shot’ Enhancements of Electron Microscopy Images of SARS-CoV-1 viruses using SSSC trained on 9 × 9 EM image patches (see Methods for details). The noisy input was obtained based on a freely available image by Gelderblom et al.. We converted the image provided by Gelderblom et al. to grayscale, cropped a section showing four viruses and scaled the resulting image to a size of 379 × 305 pixels using the same procedure as described above (see Results for details).

## Methods

The foundation of the investigated EM image enhancement approach is learning data representations from EM images of SARS-CoV-2 viruses using probabilistic generative models. Here, we apply spike-and-slab sparse coding (SSSC; Sheikh et al., 2014; Drefs et al., 2020) and a Gamma-Poisson Mixture model (GPMM; Monk et al., 2018) as examples of latent variable models with single and multiple causes. SSSC is a linear sparse coding model with binary-continuous latents and continuous, Gaussian-distributed observables:

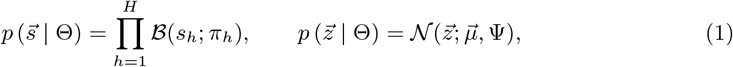

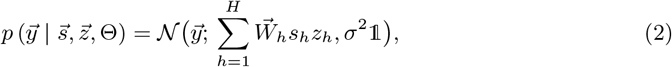

with 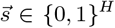, 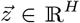 and 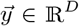 denoting latent and observed variables of the model, respectively. The binary latents follow a Bernoulli distribution 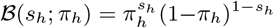 with activation probabilities *p*(*s*_*h*_ = 1) = *π*_*h*_ and *π*_*h*_ ∈ [0, 1]. The continuous latents are Gaussian-distributed with mean 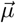 and covariance Ψ. Binary and continuous latents are combined by a point-wise multiplication, such that the resulting value has a spike-and-slab distribution, i.e. it is either equal to zero if *s*_*h*_ = 0 or continuously (normal) distributed otherwise. Given the latents, the observables follow a Gaussian whose mean is determined by a linear combination of vectors 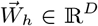 (also referred to as generative fields) which are weighted by the value of the binary-continuous latents. GPMMs, unlike SSSC, have discrete, i.e. Poisson distributed, observables 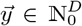 and, as mixture models, assume data points to originate from a single cause, i.e. a particular mixture component. Unlike standard Poisson mixture models, GPMMs use a continuous, Gamma-distributed latent variable 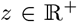 to model mixture components’ intensities. The model is defined as:

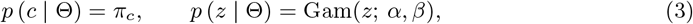

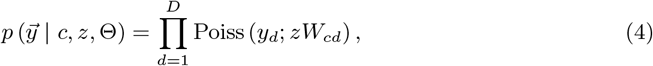

with *c* = 1 , … , *C* denoting the index of the mixture components, *π*_*c*_ ∈ [0, 1] and 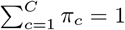, and *α, β*, 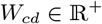.

Rather than fitting models to entire EM images, we segment a given image into overlapping patches (compare, e.g., Burger et al., 2012; Sheikh et al., 2014; Drefs et al., 2020) and use these as the data set based on which we train SSSC models and GPMMs. Given a set of data points 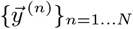, i.e., a set of EM image patches, we seek parameters Θ* that optimize the data log-likelihood:

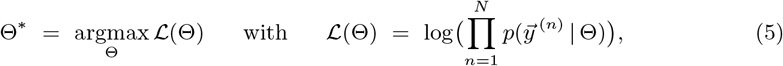

where for SSSC

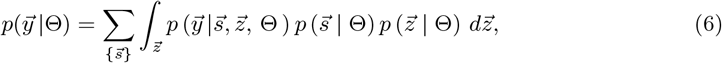

with the sum in (6) running over all possible configurations of the binary latent vector 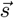, and for GPMMs

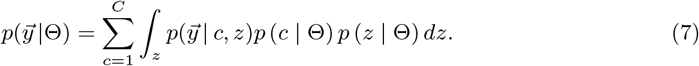

The likelihood is usually difficult to optimize directly, we therefore instead apply a variational Expectation Maximization approach (Saul and Jordan, 1996; Neal and Hinton, 1998) and optimize a variational lower bound 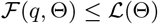 of the log-likelihood (the variational lower bound is also known as free energy or evidence lower bound). 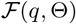 is optimized alternately w.r.t. the so-called variational distributions *q* and the model parameters Θ. As implied by the inequality, changes of the model parameters that lead to increases of the free energy also lead to increases of the log-likelihood. For GPMMs, we leverage theoretical results from Monk et al.(2018) and replace *q* by the exact posterior distribution of the model. For SSSC, on the other hand, computing exact posteriors is intractable and we instead apply a variational Expectation Maximization algorithm based on truncated posteriors (compare Drefs et al., 2020, and also see Sheikh et al., 2014).

Having inferred optimal model parameters, we can use the learned data representation in order to reconstruct the dataset, i.e. the given set of EM image patches. To this end, we apply a probabilistic data estimator derived based on the posterior predictive distribution 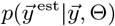.

For the SSSC model, this estimator is given by (see Drefs et al., 2020, for details):

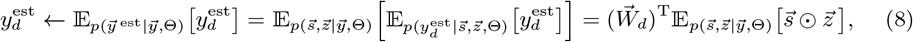

with 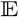 denoting expectation, 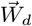 a vector containing the *d*-th entries of the generative fields 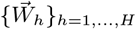 and denoting point-wise multiplication. The estimator (8) can efficiently be computed by approximating expectations w.r.t. exact posteriors by expectations w.r.t truncated posteriors (see Drefs et al., 2020, for details). We follow the same line of reasoning in order to derive a data estimator for GPMMs; the final expression takes the following form:

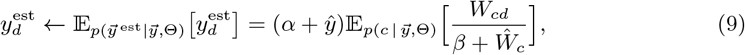

where 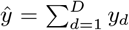 and 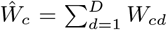. After applying (8) and (9) to each data point, i.e. to each EM image patch, we can generate a reconstructed EM image by computing means and variances of the pixel estimates based on different patches (compare Fig. 2).

## Discussion

When being produced at the limit of the possible resolution, electron microscopy (EM) images are typically contaminated by strong noise. If only noisy images are available, current mainstream approaches for denoising such as feed-forward neural networks (DNNs) can not be used because they require large datasets of pairs of clean and noisy images (e.g., Zhang et al., 2017; Tai et al., 2017; Dong et al., 2019; Tian et al., 2020). For high resolution EM images, alternative approaches are consequently required.

Here, we investigated probabilistic generative approaches which can directly learn representations from noisy images. Such approaches are by definition unsupervised and typically require much fewer data to learn appropriate representations. For the enhancement of EM images as considered here, noisy data in the form of patches extracted from a single EM image of an infection scene proved sufficient. For standard image denoising and inpainting benchmarks, recent work (Guiraud et al., 2020; Drefs et al., 2020) has shown that in the ‘zero-shot’ setting, generative approaches can represent state-of-the-art algorithms, i.e., they can achieve higher peak-signal-to-noise-ratios (PSNRs) than recent alternatives such as Noise2Void (Krull et al., 2019) or Deep Image Prior (Ulyanov et al., 2018). Visual inspection of the here shown denoised EM images confirms the benchmarking results of previous work, although a standard benchmarking is not available for the task here considered: to compute PSNR values for comparison, clean ground-truth data would be required. However, visual inspection of the here obtained de-noising results (for instance Fig. 2) strongly suggests that the competitive performance carries over and shows effectively suppressing noise in high-resolution EM recordings.

In addition to the estimation of denoised images, we have here investigated higher-order reconstruction statistics. Using the variances of pixel estimations, images can be computed which recover a spatial impression of the corresponding infection scene. As for the conventionally denoised images, we computed the spatially appearing images based on a single, noisy TEM image of the scene. This stands in contrast to the procedure that Nanographics (2021) have applied to produce 3D reconstructions of a single SARS-CoV-2 virus. On the one hand, such a tomogram reconstruction of the virus is more detailed and can nicely visualize the spike protein structure and distribution on the virus’ surface. On the other hand, the method relies on many TEM images of the same single virus, and many images are more difficult to obtain than single images. Furthermore, whole infection scenes are more challenging to reconstruct than a single virus, and 3D tomogram reconstructions of whole such scenes have (to our knowledge) not been reported so far. Importantly, different approaches for 3D reconstruction come with different artifacts. Tomogram approaches can misestimate surfaces especially for noisy data, while our reconstructions based on generative models can introduce artifacts, e.g., because rare structures are underrepresented. Furthermore, due to being single-image based, actual surfaces are not estimated by our approach. An advantage is that surface structures do not occlude other structures – the here shown images have a ‘glassy’ effect. A disadvantage is that, e.g., internal structures of the virus can get entangled with surface structures. To identify single spikes across the whole surface of the virus, the 3D reconstruction method of Nanographics (2021) is consequently preferable. In general, however, as artifacts are different for the different approaches, the methods can be used to mutually confirm the true nano-scale structures that are visualized. Furthermore, different methods can potentially be combined in hybrid systems. For instance, ‘zero-shot’ denoising of single images by generative models could be used to generate denoised images for improved 3D surface reconstruction with methods used by Nanographics (2021); or ‘zero-shot’ denoising could be used in conjunction with other EM imaging methods (e.g. Titze and Genoud, 2016). Finally, different approaches are applicable to different types of available data. The method for protein reconstruction (e.g. Wrapp et al., 2020; Ke et al., 2020) combines TEM images of many protein molecules of many different viruses, which neither the tomogram method of Nanographics (2021) nor our approach is capable of. Also, unlike the surface reconstruction by Nanographics (2021), our approach can not combine many TEM images of the same virus to generate a 3D tomogram. However, neither the method of Wrapp et al. (2020); Ke et al. (2020) nor of Nanographics (2021) can be used to recover a spatial impression from a single noisy TEM image. The here presented method is consequently able to recover images giving a spatial impression of whole infection scenes that can enhance small details. Images showing virus infections spatially have been made available previously, e.g., by using scanning electron microscopes (see e.g. NIAID, 2020, for examples). Such spatial images are at a lower resolution, however, and do therefore not show the characteristic details of SARS-CoV-2 viruses, for instance. In contrast, we here show images the give a spatial impression of whole infection scenes at very high resolution. To our knowledge, these are the first images at such a level of detail and the first for a SARS-CoV-2 infection scene. Future work could, for instance, compare different EM recordings of the same images of SARS-CoV-2 infection scenes (as well as other scenes). Such comparisons would help in better understanding the ‘glassy’ effect we observe, and how internal and external structures of the virus get entangeled using our approach.

In general, beyond their ability to provide more details, we believe that the here shown images with their easy to identify corona viruses can more plastically and more directly communicate the threat posed by the virus. In turn, it may contribute to the motivation of each individual to help battling the COVID-19 pandemic.

## Acknowledgements

We would like to acknowledge funding by the German Ministry of Research and Education (BMBF) in the project 05M2020 (SPAplus) which enabled this research through a top-up fund for COVID-19 research; and we would like to acknowledge funding by the DFG project 352015383 (SFB 1330, B2) which provided source code to train generative models. Furthermore, we would like to acknowledge support in terms of computational resources by the Oldenburg High Performance Compute Cluster (CARL) and by the North German Supercomputing Alliance under grant nim00006.

