## Supplementary figures and images for "Visualization of SARS-CoV-2 Infection Scenes by ‘Zero-Shot’ Enhancements of Electron Microscopy Images"

### Supplementary Figure 1

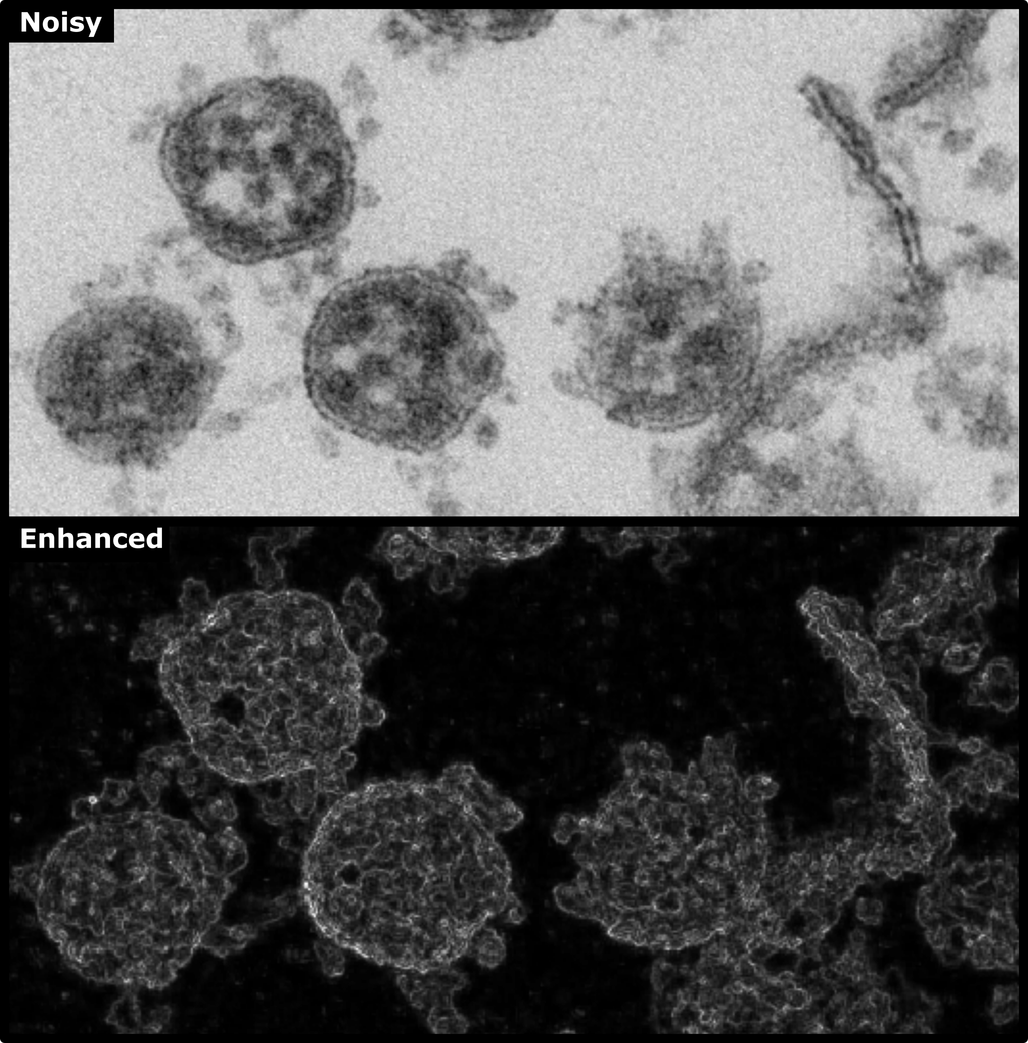
